# Multifaceted role of RIMBP2 in promoting hearing in murine cochlear hair cells

**DOI:** 10.1101/2022.05.03.490398

**Authors:** Menghui Liao, Xin Chen, Ling Lu, Rongrong Guo, Panpan Zhang, Yige Li, Yuhua Zhang, Qiaojun Fang, Yangnan Hu, Jiaying Cai, Xiaoyan Chen, Mingliang Tang, Xia Gao, Shuijin He, Huawei Li, Geng-Lin Li, Renjie Chai

**Author notes:** Corresponding authors: Drs. Renjie Chai, Geng-Lin Li, and Huawei Li. These authors contributed equally to this paper (Menghui Liao, Xin Chen, Ling Lu).

## Abstract

In peripheral, the mammalian cochlea is a remarkable sensory apparatus, owning to its outer and inner hair cells (OHCs and IHCs) that amplify and transmit auditory signals to the brain, respectively. Rab3-interacting molecular (RIM) binding protein 2 (RIMBP2) is widely expressed in receptor cells and neurons, specifically in the active zones for exocytosis of synaptic vesicles (SVs), but its exact functions in the cochlea are not very well understood. We therefore generated a *Rimbp2* knockout mouse model (*Rimbp2-/-*), which exhibited severely impaired hearing with not only elevated hearing thresholds but also increased latencies and reduced amplitudes in the Wave I of their auditory brainstem responses (ABRs). Consistent with the threshold elevation, we found significant loss of OHCs, likely through apoptosis, in the *Rimbp2-/-* cochlea. Consistent with changes observed in the ABR Wave I, we found greatly reduced exocytosis, both spontaneously and upon stimulation, in *Rimbp2-/-* IHCs. Specifically, our patch-clamp analysis on IHCs and postsynaptic spiral ganglion neurons (SGNs) revealed not only reduced readily releasable pool (RRP) of SVs but also reduced sustained release rate, along with complete blockade of fast endocytosis, in *Rimbp2-/-* IHCs. Lastly, with immunostaining of whole-mounted cochleae, we found that while the number of ribbon synapses in *Rimbp2-/-* IHCs was unchanged, their localization moved subtly but significantly towards the basal pole of IHCs. Taken together, we uncovered an unexpected role of RIMBP2 for OHC survival and a more extensive role in promoting IHC exocytosis than previously believed.

## Introduction

In mammals, hearing starts in the cochlea in that it decomposes acoustic signals into different frequencies, which vibrate the organ of Corti at designated locations along the winding cochlear duct (Fettiplace & Hackney, 2006). At each location, there are three rows of OHCs and one row of IHCs, both of which are equipped with stereocilia at their apical pole with mechanoelectrical transduction (MET) channels to convert mechanical vibrations into membrane potential changes but produce distinct outputs. OHCs employ densely packed prestin, a motor protein, in their lateral membrane to amplify vibration, conferring the cochlea with remarkable sensitivity of low level sounds (Fettiplace & Hackney, 2006). It has been shown that OHCs are extremely vulnerable to noise, genetic mutations, drugs and aging (Wagner & Shin, 2019), and loss of OHCs causes significant elevation of hearing thresholds (Liberman & Dodds, 1984). IHCs, on the other hand, are mainly responsible for sending sensory signals to the brain, through ribbon synapses with axonal boutons of spiral ganglion neurons (SGNs) in their basal pole (Rutherford & Pangrsic, 2012). These ribbon synapses are specialized with multivesicular release (Glowatzki E, 2002; Li, Keen, Andor-Ardo, Hudspeth, & von Gersdorff, 2009), large readily releasable pool (RRP) of synaptic vesicles (SVs), and tight coupling of Ca^2+^ channels and docked SVs (Goutman & Glowatzki, 2007; Pangrsic et al., 2015), allowing them to operate with extraordinary temporal precision.

At synapses of both ribbon and non-ribbon types, fusion of SVs occurs at the active zones, which contain sophisticated cytomatrix to coordinate the process in a well- controlled manner (Sudhof, 2012). Among a host of proteins that make up the cytomatrix in the active zones, Rab3-interacting molecular (RIM) binding protein 2 (RIMBP2) is widely believed to play a central role (Mittelstaedt & Schoch, 2007). As the other two members of the highly conserved RIMBP family, RIMBP2 has three SH3 domains, one in the middle and two at the C-terminal, allowing it to bind to not only RIMs, which is an integral player itself in the cytomatrix of the active zones, but also voltage-gated Ca^2+^ channels for Ca^2+^ influx that triggers fusion of SVs (Hibino et al., 2002; Mittelstaedt & Schoch, 2007). Unsurprisingly, deletion of *Rimbp2* bears significant impact on synaptic transmission in a range of synapses in different species. In Drosophila neuromuscular junction (NMJ), it reduces Ca^2+^ channel abundance and Ca^2+^ influx, loosens coupling of Ca^2+^ channels and SVs, decreases the size of RRP, and slows replenishment of SVs, rendering a total of ∼ 10-fold decrease in synaptic transmission (Liu et al., 2011; Muller, Genc, & Davis, 2015). At murine hippocampal synapses, deletion of *Rimbp2* reduces Ca^2+^ clustering without affecting Ca^2+^ channel abundance or Ca^2+^ influx, and loosens coupling of Ca^2+^ channels and SVs without changing the size of RRP or replenishment of SVs (Grauel et al., 2016). At the calyx of Held synapses in the auditory brainstem, deletion of *Rimbp2* fails to change the size of RRP as well, but it alters the dynamics of its release, which combines with impaired coupling of Ca^2+^ channels and SVs and makes the synapses significantly less reliable (Acuna, Liu, Gonzalez, & Sudhof, 2015). At the endbulb of Held synapses, also in the auditory brainstem, deletion of *Rimbp2* reduces SV tethering and docking, and therefore slows replenishment of SVs (Butola et al., 2021).

In ribbon synapses, deletion of *Rimbp2* also caused significant changes in synaptic transmission. In the retinal ribbon synapse between rod bipolar cell and AII amacrine cell, double knockout of *Rimbp2* and *Rimbp1* reduced both Ca^2+^ channel density and Ca^2+^ current, loosened the coupling of Ca^2+^ channels and SVs, decreased and desynchronized exocytosis, and impaired both emptying and replenishment of RRP without changing its size (Luo, Liu, Sudhof, & Acuna, 2017). In ribbon synapses of IHCs, deletion of *Rimbp2* also reduced Ca^2+^ influx and exocytosis, but the coupling between the two appeared be unchanged (Krinner, Butola, Jung, Wichmann, & Moser, 2017). In this same study, while the evidence for reduced replenishment of SVs was convincing, the evidence for RRP showed a trend of reduction but failed to reach significance (Krinner et al., 2017), leaving the matter remained to be carefully examined. Furthermore, IHCs release SVs spontaneously at a high rate, both single and multivesicular (Glowatzki E, 2002; Li et al., 2009), and it is unknown if deletion of *Rimbp2* bears any impact on this hallmark phenomenon in IHCs. To address these open questions, we generated a new *Rimbp2* knockout mouse model, which, unlike the previous one with only mild hearing loss (Krinner et al., 2017; Krinner, Predoehl, Burfeind, Vogl, & Moser, 2021), exhibited severe hearing impairment. Furthermore, in the cochlea of this new mouse model, we found not only significant loss of OHCs but also extensive changes of exocytosis in IHCs, therefore revealing a multifaceted role of *Rimbp2* in promoting hearing.

## Results

### Expression of RIMBP2 in the mouse cochlea

To start, we sought out to examine the expression of RIMBP2 in cochlea. We chose postnatal day 0 (P0) wild-type (WT) mice for immunolabeling RIMBP2 in the cochlear epithelia together with Myosin7a and Sox2, which label specifically hair cells (HCs) and supporting cells (SCs), respectively. Through confocal images of the whole-mounted organ of Corti, it was observed that the majority of RIMBP2 was present in the cytoplasm of HCs at P0 (Fig. 1a). In frozen cochlear sections, immunofluorescence staining confirmed that RIMBP2 was located in HCs (Fig. 1b), including both outer and inner hair cells (OHCs and IHCs). We also examined RIMBP2 expression for late developmental stages, and we found similar expression pattern from P14 to P30 (Supplementary Fig. 1). No significant difference was found in RIMBP2 labelling in the three turns. To further verify the expression of *Rimbp2* gene, we performed western blot and q-PCR analyses with the WT mouse cochlea. Western blot showed that a specific band of RIMBP2 (116 kDa), was detected in both the cochlear and the brain from WT mice (Fig. 1c). Furthermore, q-PCR analysis revealed that from E16 to P30 *Rimbp2* was expressed at the mRNA level in the mouse cochlea (Fig. 1d). Together, these results indicate that RIMBP2 was expressed in the mouse cochlea continuously from embryonic to adult stage.

**Figure 1.**
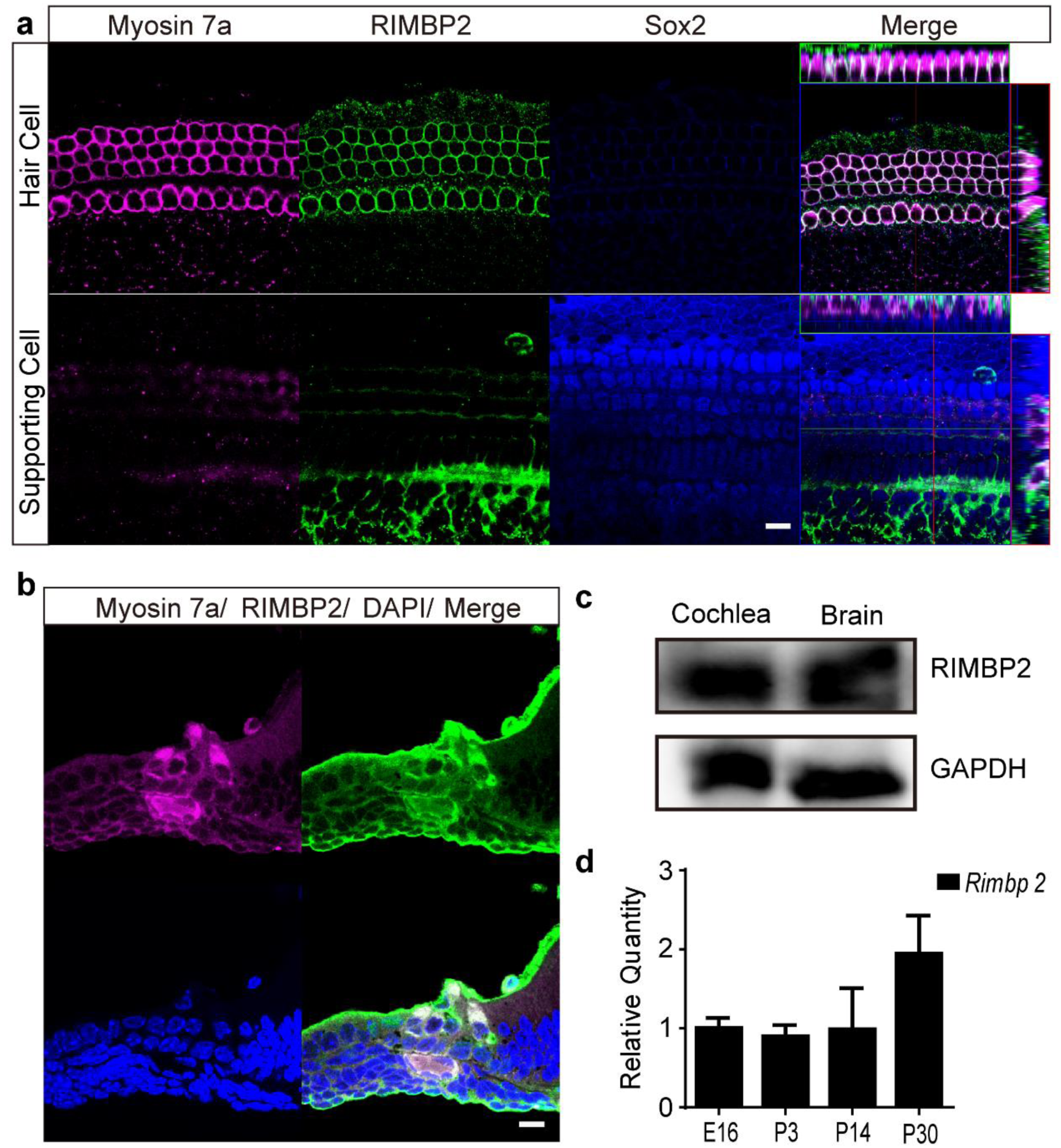
Expression of RIMBP2 in the mouse cochlea. (a) Immunofluorescence staining of whole-mounted cochleae (P0) showing that RIMBP2 (green) was expressed in both OHCs and IHCs in the mouse cochlea. Images showed the middle turn of the cochlear basement membrane. Myosin7a was used as a marker for hair cells (magenta) and Sox2 for supporting cells (blue). (b) Immunofluorescence staining of frozen cochlear sections (P0) showing that RIMBP2 was expressed in OHCs and IHCs. (c) Western blot analysis of the mouse cochlea and brain at P0. (d) Expression of RMBP2 at the mRNA level in the mouse cochlea over development. The internal control for both c and d was GAPDH. Data are presented as mean ± SD. Scale bars = 10 μm. Figure 1-source data 1 Related to Figure 1c. Figure 1-source data 2 Related to Figure 1d.

### Generation of Rimbp2 knockout mice

We demonstrated that *Rimbp2* was consistently expressed in HCs in the mouse cochlea. So, what are the possible functions for it to have on hearing? To address this question, we constructed a *Rimbp2* knockout (*Rimbp2-/-*) mouse model through the PB Transposon system (Sheng Ding 2005). To verify whether the gene was successfully knocked out, we sequenced the PCR product, and we found that a 791-bp sequence was successfully inserted in the *Rimbp2* gene, before TTAA, as planned (Fig. 2a). We then performed Western blot analysis and found that the specific band was detected in the cochlear from WT mice, but not the *Rimbp2-/-* mice (Fig. 2b). Furthermore, q-PCR result confirmed that the *Rimbp2* gene was knocked out nearly completely in the *Rimbp2-/-* mice (WT: 1.00±0.09; *Rimbp2*-/-: 0.01±0.00; p = 0.0028, Fig. 2c, the primer sequences are listed in Supplementary Table 2). Lastly, the immunolabeling of RIMBP2 was clearly seen in cochlear HCs of WT mice, while it was hardly found in the *Rimbp2-/-* cochlea (Fig. 2d). The weak signal of immunolabeling in the *Rimbp2-/-* cochlea was likely due to the non-specific binding of antibodies. Taken all these results together, we conclude that the *Rimbp2* gene had been successfully knocked out in the *Rimbp2-/-* mouse model.

**Figure 2.**
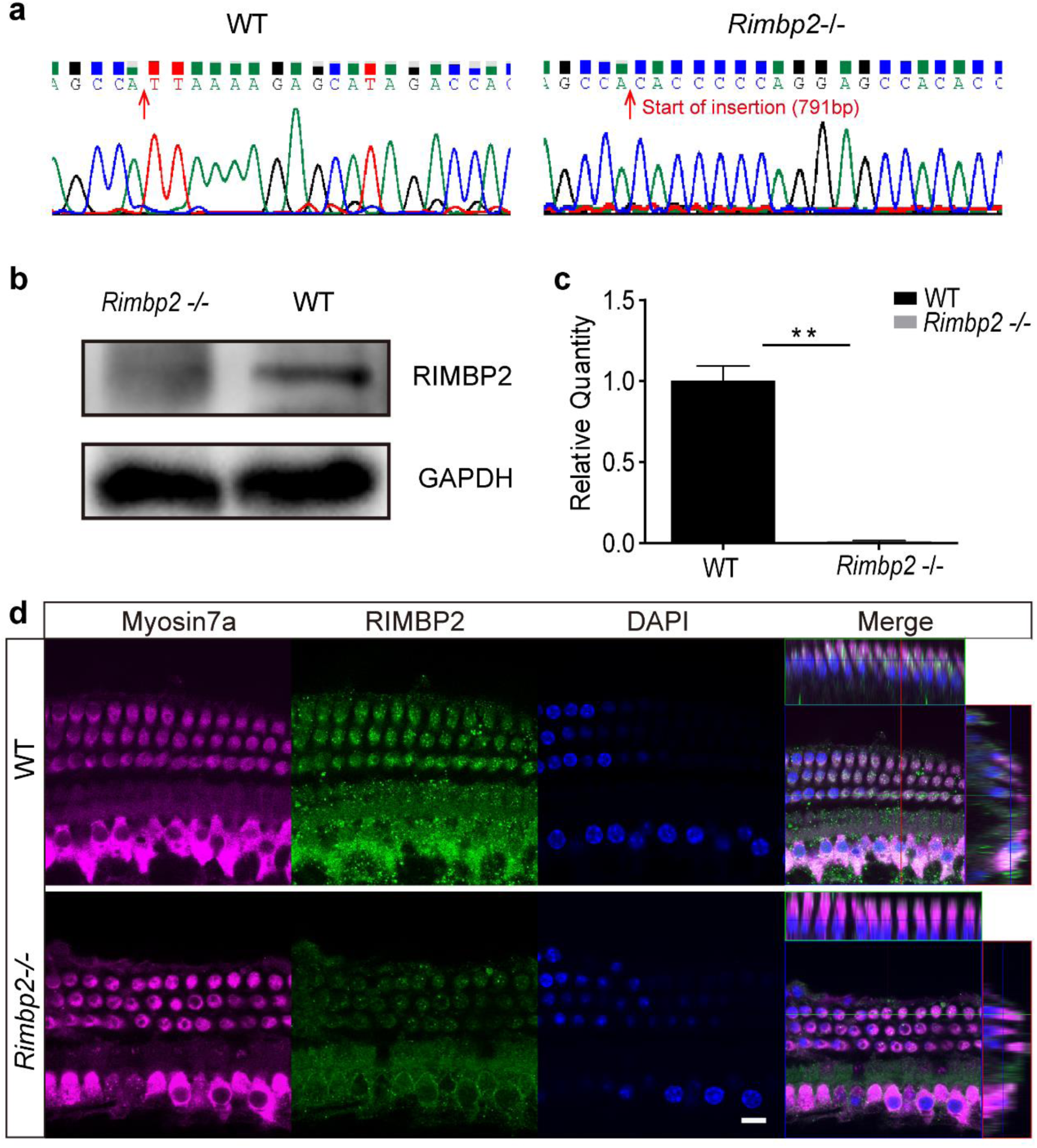
Generation of *Rimbp2* knockout mice. (a) Sequencing of PCR products from the cochlea of WT and *Rimbp2-/-* mice. (b) Western blot analysis of RIMBP2 in WT and the *Rimbp2-/-* cochlea at P0. (c) *Rimbp2* RNA level in the cochlea of WT and *Rimbp2-/-* mice at P0 (**: p<0.01). (d) Immunofluorescence images of the WT and *Rimbp2-/-* cochleae at P21, showing elimination of RIMBP2 at the protein level. Data are presented as mean ± SD. Scale bar = 10 μm. Figure 2-source data 1 Related to Figure 2b. Figure 2-source data 2 Related to Figure 2c.

### Impaired hearing of Rimbp2-/- mice

We next set out to investigate possible changes in hearing of *Rimbp2-/-* mice. Given that the auditory brainstem response (ABR) reflects the overall functionality of the cochlear output (Felicia Gilels 2017), we therefore recorded ABRs in response to pure tone pips in WT and *Rimbp2-/-* mice under anesthesia. We found that the deletion of *Rimbp2* caused severe hearing loss, in range of 20 - 50 dB, significantly more severe than previously reported with a different mouse model (Krinner et al., 2017; Krinner et al., 2021), across all frequencies tested, and in all three ages tested (Fig. 3a, see Supplementary Table 3-1 for numbers). For further analysis, we chose ABR recordings obtained from WT and *Rimbp2-/-* mice at P21. As shown in Fig. 3b obtained from a pair of animals, while the WT mouse responded to sound stimulation at low as 30 dB, no ABR waveform can be discernable for sound below 75 dB in the *Rimbp2-/-* mouse. To shed light on the underlying mechanisms of this hearing loss caused by *Rimbp2* deletion, we examined Wave I latency and amplitude for sound stimulation of 8 and 16 kHz. As shown in Fig. 3c, we found that at 80 dB, the Wave I latency was significantly increased from 1.93±0.11 and 1.86±0.19 ms to 2.31±0.12 and 2.32±0.26 ms, for 8 and 16 kHz, respectively, suggesting changes in OHCs in the cochlea of *Rimbp2-/-* mice. Meanwhile, at the same sound pressure level, the Wave I amplitude was significantly reduced from 0.58±0.09 and 0.88±0.11 μV to 0.24±0.21 and 0.23±0.19 μV (Fig. 3d), for 8 and 16 kHz, respectively, indicating changes in IHCs in the cochlea of *Rimbp2-/-* mice. Together, these results suggest that the hearing impairment caused by *Rimbp2* deletion is likely due to changes in both OHCs and IHCs in the mouse cochlea.

**Figure 3.**
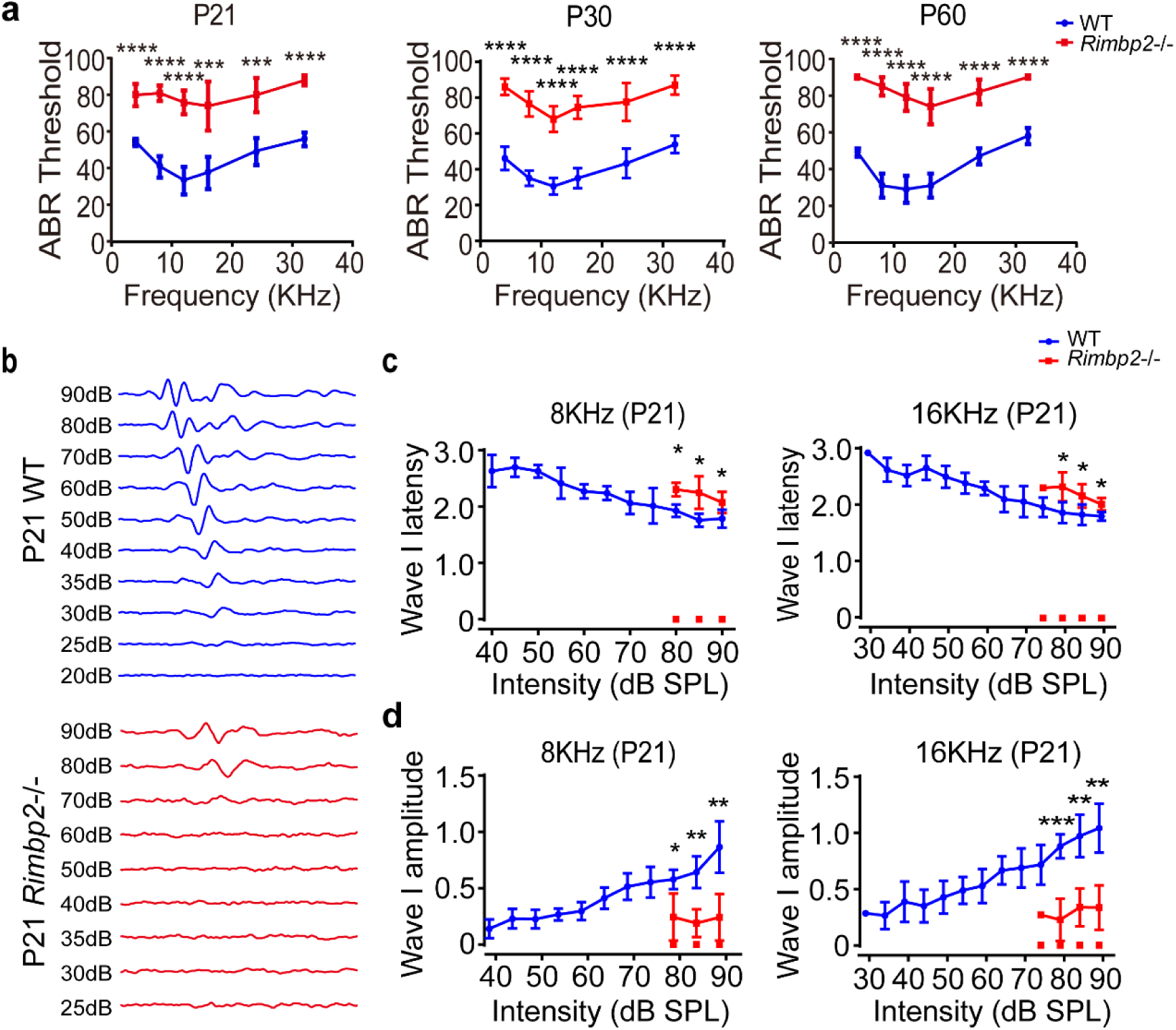
Hearing of *Rimbp2*-/- mice was severely impaired. (a) ABR thresholds for WT (blue) and *Rimbp2-/-* mice (red) at P21, P30 and P60. ABR thresholds were assessed at 4, 8, 12, 16, 24, and 32 kHz with pure tone pips. (b) ABR waveforms of a WT and *Rimbp2-/-* mouse pair at P21 for tone pips of 16 kHz. (c) ABR wave Ⅰ latency of WT and *Rimbp2-/-* mice at P21 for tone pips of 8 and 16 kHz, showing the latency was greatly increased in KO mice. (d) ABR wave Ⅰ amplitude of WT and *Rimbp2-/-* mice at P21 for tone pips of 8 and 16 kHz, showing the amplitude was greatly decreased in KO mice. Data are presented as mean ± SD (*: p<0.05, **: p<0.01, ***: p<0.001, ****: p<0.0001). Figure 3-source data 1 Related to Figure 3a. Figure 3-source data 2 Related to Figure 3c and d.

### Apoptosis of OHCs in Rimbp2-/- mice

The ABR results described above on the *Rimpb2*-/- mice led us to consider changes in both OHCs and IHCs. We therefore fixed cochleae from the *Rimpb2* KO mice and their littermates at P21, P30 and P60, and performed immunofluorescence staining on the whole-mounted organ of Corti. While IHCs appeared to be intact, there were void spaces where OHCs were missing, suggesting loss of OHCs (Fig. 4a-c). We counted both types of hair cells, and we found significant loss of OHCs (P21, in the apical turn, WT: 39.6±1.70 per 100 μm, *Rimbp2-/-*: 37.2±2.94 per 100 μm), but not IHCs (P21, in the apical turn, WT: 12.3±0.49 per 100 μm, and *Rimbp2-/-*: 12.1±0.61 per 100 μm; Fig. 4d-i, Supplementary Table 4). We went on to examine the stereocilia of both types of HCs in the *Rimbp2* KO cochlea, and we found that both of its morphology and arrangement appeared normal (Supplementary Fig. 2).

**Figure 4.**
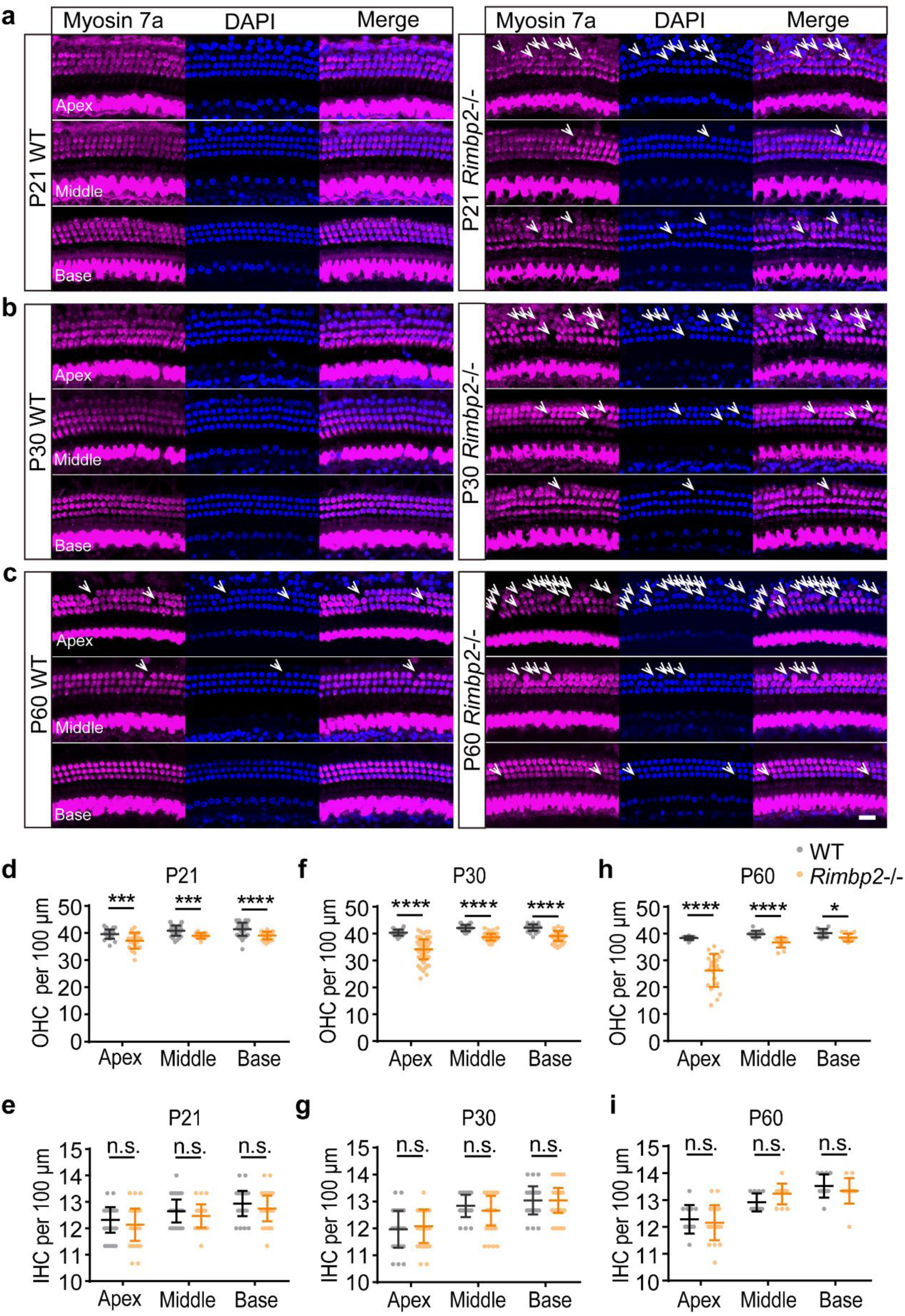
Knockout of *Rimbp2* caused significant loss of cochlear OHCs but not IHCs. (a-c) Immunofluorescence images of WT and *Rimbp2-/-* cochleae at P21 (a), P30 (b) and P60 (c). Hair cells were stained with anti-Myosin7a antibody (magenta). Images were taken from the apical, middle and basal turn of cochlea. Arrows indicate missing outer hair cells. (d-i) Hair cell densities in the cochlea of WT (gray) and *Rimbp2-/-* mice (yellow) at P21, P30 and P60, showing significant loss of OHCs (d, f, h), but not IHCs (e, g, i). Data are presented as mean ± SD (*: p<0.05, ***: p<0.001, ****: p<0.0001). Scale bar = 20 μm. Figure 4-source data 1 Related to Figure 4d, f and h.

Given the significant loss of OHCs in the *Rimbp2-/-* cochlea, we hypothesized that it was caused by apoptosis, and we therefore decided to examine expression of TUNEL, a key apoptotic factor, in the cochlea at an earlier age. We collected cochleae from *Rimbp2-/-* mice and their littermate WT mice at P18 and performed double immunofluorescence staining against TUNEL and MYOSIN7a. As shown in Fig. 5a, we found significantly more double positive cells in the *Rimbp2-/-* cochlea than in the WT cochlea, for all the three turns (apical: WT: 0.00±0.00, *Rimbp2*-/-: 0.08±0.02, n = 3, p = 0.0028; middle: WT: 0.00±0.00, *Rimbp2*-/-: 0.07±0.00, n = 3, p < 0.0001; basal: WT: 0.00±0.00, *Rimbp2*-/-: 0.08±0.01, n = 3, p = 0.0006; Fig. 5b). We also performed Western blot analysis of cleaved-*Caspase3*, another key apoptotic factor, and we found its expression was observably increased in the *Rimbp2-/-* cochlea (WT: 0.59±0.08, *Rimbp2*-/-: 1.04±0.09, p = 0.0028, Fig. 5c-d). Furthermore, we examined the mRNA expression level of four other apoptosis-related factors, and we found that while *Caspase3* (WT: 1.03±0.32, *Rimbp2*-/-: 1.06±0.21, p = 0.8961) and *Bcl-2* (WT: 1.00±0.07, *Rimbp2*-/-: 1.17±0.26, p = 0.3860) did not differ significantly in the two types of cochlea, *Caspase2* (WT: 1.01±0.18, *Rimbp2*-/-: 2.62±0.18, p = 0.0004) and *Apaf1* (WT: 1.00±0.06, *Rimbp2*-/-: 2.11±0.10, p = 0.0002) were significantly increased in the cochlea of *Rimbp2-/-* mice (Fig. 5e). Taken together, these results show that *Rimbp2* deletion caused loss of OHCs in the cochlea through apoptosis, suggesting *Rimbp2* was involved in the maintenance and survival of OHCs.

**Figure 5.**
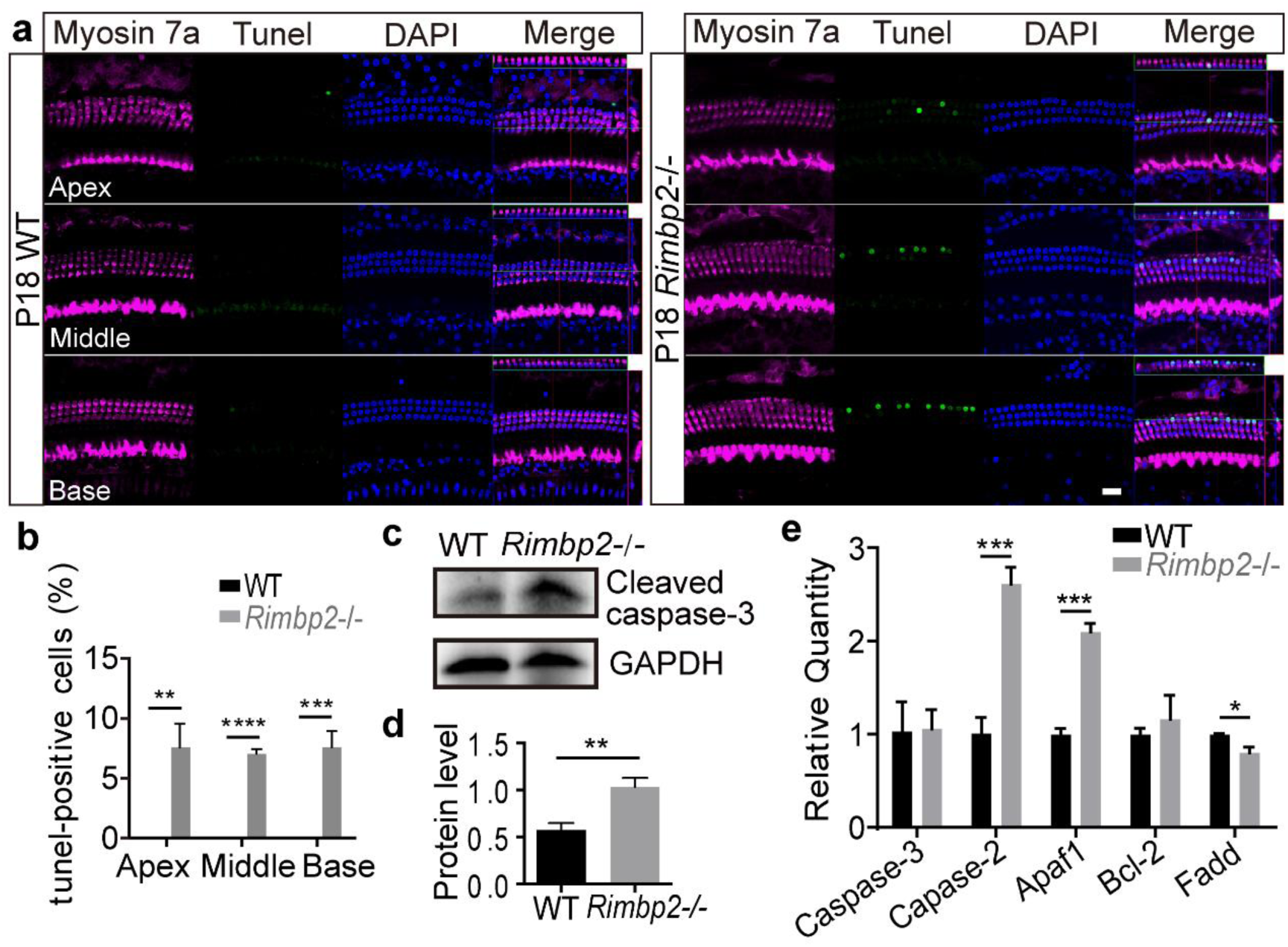
Knockout of *Rimbp2* led to apoptosis in OHCs in the developing cochlea. (a) Immunofluorescence images showing Tunel (green) and Myosin7a (magenta) double-positive cells appeared in the P18 cochlea of *Rimbp2-/-* but not WT mice. (b) Numbers of Tunel and Myosin7a double positive cells in the apical, middle and basal turn of the WT (black) and *Rimbp2-/-* cochleae (gray). (c-d) Western blot analysis showing that the expression level of Cleaved-Caspase3 was significantly increased in the *Rimbp2-/-* cochlea. (e) RNA expression levels of apoptosis regulators in the WT and *Rimbp2-/-* cochleae. Data are presented as mean ± SD (*: p<0.05, **: p<0.01, ***: p<0.001, ****: p<0.0001). Scale bar = 20 μm. Figure 5-source data 1 Related to Figure 5b. Figure 5-source data 2 Related to Figure 5c. Figure 5-source data 3 Related to Figure 5d. Figure 5-source data 4 Related to Figure 5e.

### Reduced exocytosis in IHCs in Rimbp2-/- mice

Our ABR analysis of impaired hearing in *Rimbp2*-/- mice indicated changes in both OHCs and IHCs (Fig. 3), but we found loss of OHCs only, not IHCs (Fig. 4). To examine possible functional changes in IHCs, we performed patch-clamp recording to study their exocytosis, a key step allowing them to transmit sensory signals to the brain through SGNs. Under voltage-clamp, we applied voltage steps of increasing duration from 2 to 500 ms, from a holding potential of -90 mV to 0 mV, which induced a L-type Ca^2+^ current (I_Ca_) and an increase of whole-cell capacitance (ΔC_m_) indicating associated exocytosis (Fig. 6a). We found no significant change in I_Ca_, as indicated by Ca^2+^ charge, i.e. the integral of I_Ca_ over time, for all stimulation durations tested (Fig. 6b, left). For ΔC_m_, there was no significant difference between WT and *Rimbp2*-/- IHCs, either, for stimulation duration of 2 to 200 ms, but for stimulation of 500 ms, ΔC_m_ was markedly reduced in *Rimbp2*-/- IHCs (WT: 73.7±30.4 fF, *Rimbp2-/-*: 45.7±8.44 fF, p = 0.0111, Fig. 6b, right), suggesting that *Rimbp2* deletion reduced sustained exocytosis in IHCs. To further examine the impact of *Rimbp2* deletion on dynamics of exocytosis in IHCs, we plotted ΔC_m_ against the stimulation duration and fitted the data to a simple release and refilling model of exocytosis (Fig. 6c), which yielded a RRP of SVs, a time constant to release RRP and a sustained release rate (see Materials and Methods). In addition, given the negative values of ΔC_m_ for stimulation of 2 and 5 ms in WT mice, which turned positive in *Rimbp2* knockout mice (Fig. 6b, right, inset), we suspected that there was fast endocytosis that was eliminated by *Rimbp2* deletion, we therefore added a constant as the fourth parameter to account for this fast endocytosis (Fig. 6c). For the time constant to release RRP, there was no significant change. But we found significant reduction in RRP from 484±248 vesicles for WT IHCs to 261±94 vesicles in *Rimbp2-/-* IHCs, and significant reduction in the sustained release rate from 3474±1476 vesicles/s to 2080±750 vesicles/s, indicating an active role of *Rimbp2* in promoting exocytosis in IHCs. Furthermore, we found the fast endocytosis was almost completely eliminated from 260±102 vesicles for WT IHCs to -5±131 vesicles for *Rimbp2* IHCs (Fig. 6c), revealing the critical role of *Rimbp2* in regulating endocytosis in IHCs.

**Figure 6.**
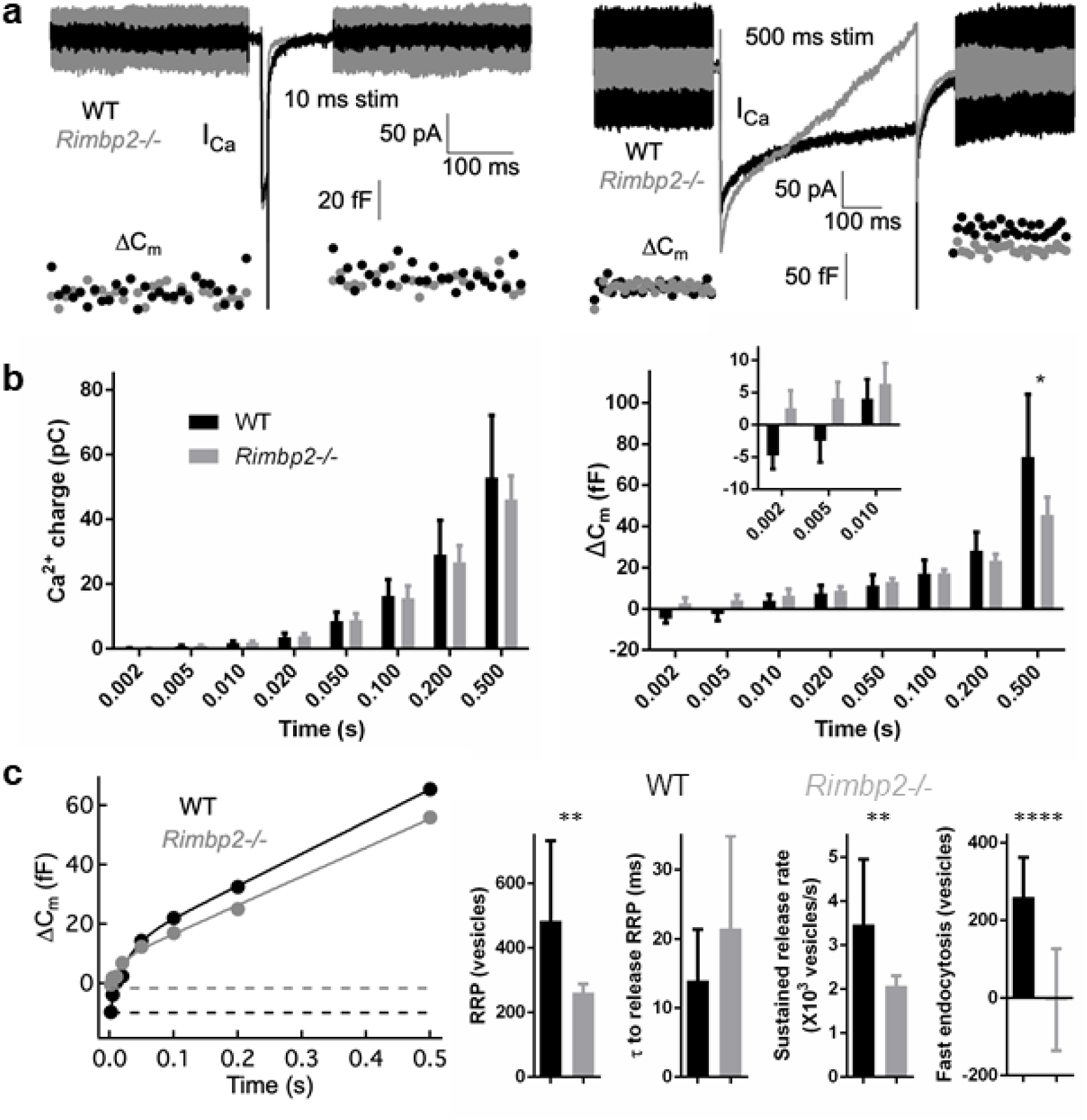
Knockout of *Rimbp2* reduced evoked exocytosis in IHCs of the mouse cochlea. (a) Ca^2+^ current (I_Ca_) and capacitance increase (ΔC_m_) induced in a pair of WT (black) and *Rimbp2-/-* IHCs (gray) in response to voltage steps of 10 (left) and 500 ms (right). ΔC_m_ was reduced for 500 ms but not for 10 ms. (b) Pooled data for Ca^2+^ charge and ΔC_m_ for all stimulation durations assessed. No significant change was observed for Ca^2+^ charge (left), and ΔC_m_ was significantly reduced for stimulation of 500 ms only (right). It is notable that the ΔC_m_ average was negative for 2 and 10 ms and turned positive for 10 ms in WT IHCs, while they were all positive for *Rimbp2-/-* IHCs (inset). (c) Plots of ΔC_m_ versus stimulation time for a pair of WT (black) and *Rimbp2-/-* IHCs (gray). The solid lines are fitting curves for extracting parameters of exocytosis (see Materials and Methods), yielding RRP, time constant (τ) to release RRP, sustained release rate, and fast endocytosis (indicated by dashed lines in the plot). (d) While τ to release RRP was not significantly altered, all the other three parameters were significantly reduced, suggesting exocytosis in IHCs was severely impaired in the *Rimbp2-/-* mouse cochlea. Data are presented as mean ± SD (*: p<0.05, **: p<0.01, ****: p<0.0001). Figure 6-source data 1 Related to Figure 6b and c.

The nearly complete blockade of fast endocytosis in IHCs in the *Rimbp2-/-* cochlea is likely to bear consequence on their spontaneous exocytosis, which occurs at a high rate even without sound stimulation (Glowatzki E, 2002; Li et al., 2009). Given that the capacitance measurement in IHCs is unable to resolve their spontaneous exocytosis, we turned to patch-clamp recording on their postsynaptic SGNs. For improved success rate, we performed patch-clamp recording on the SGN soma. When filled with neurobiotin, SGNs can be seen to extend a peripheral axon and make synaptic contact with a single IHC (Fig. 7a). Under current-clamp, we recorded spontaneous spikes in SGNs (Fig. 7b), and we found that the rate of spontaneous spikes was greatly reduced in the *Rimbp2-/-* cochlea (number of spikes in 5 mins, WT: 40.5±16.1, *Rimbp2*-/-: 14.0±10.1, p = 0.0112, Fig. 7c), in agreement with reduced spontaneous exocytosis from IHCs as we can expect from the blockade of fast endocytosis. Furthermore, the rate, duration and spike number of spontaneous spike bursts were almost completely diminished, consistent with the reduction of sustained release rate of exocytosis in IHCs described above.

**Figure 7.**
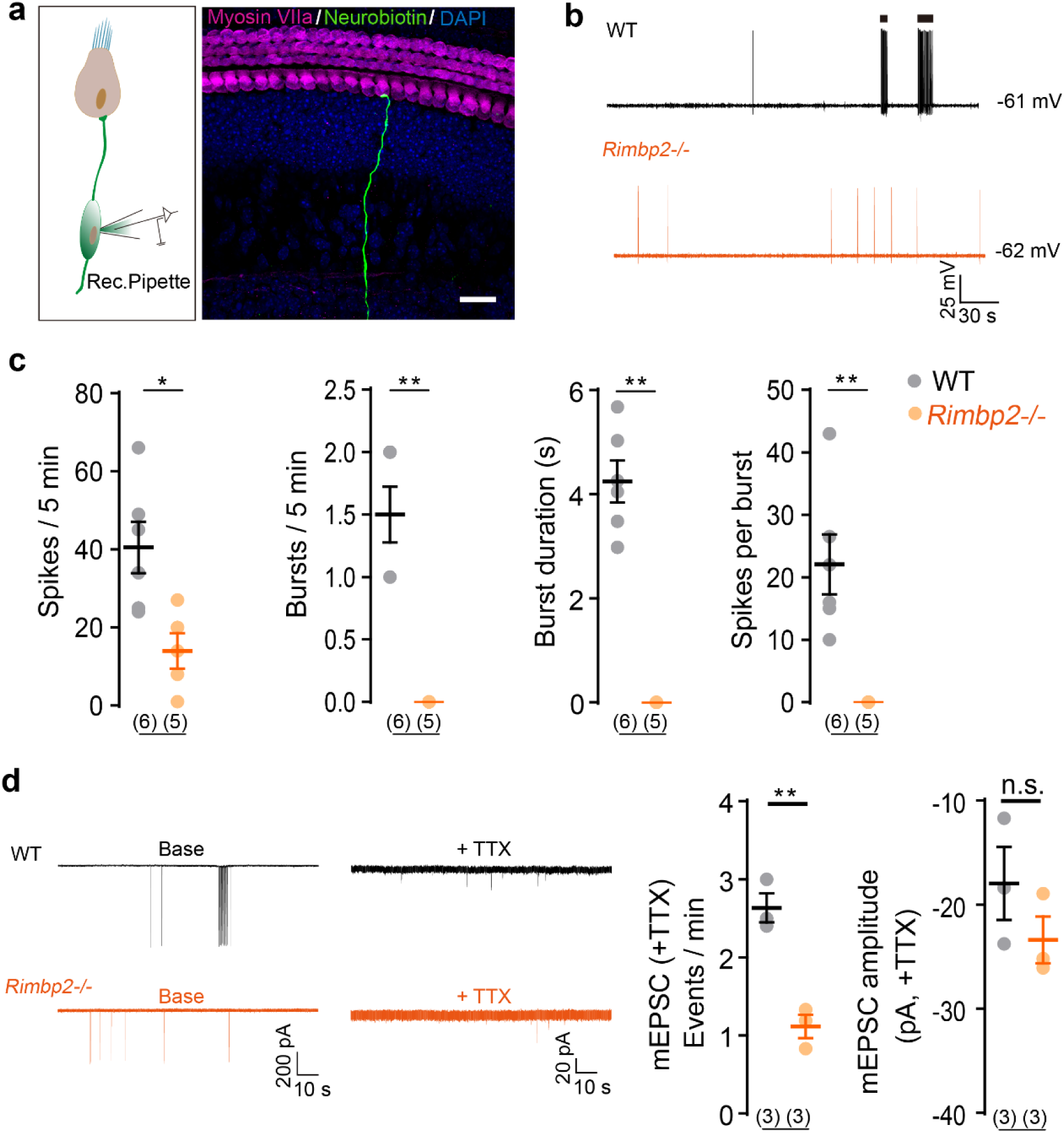
Knockout of *Rimbp2* reduced the spontaneous exocytosis in IHCs of the mouse cochlea. (a) Patch-clamp recording diagram in the soma of SGNs in the cochlea (left) and tracking of the peripheral axon and its terminal on an IHC (right). (b) Current-clamp recording in a pair of WT (black) and *Rimbp2*-/- SGNs (yellow), showing that the spontaneous spikes were greatly reduced in the *Rimbp2-/-* SGN. (c) Pooled data showing spontaneous spikes in SGNs were greatly diminished in the *Rimbp2-/-* cochlea. (d) Voltage-clamp recording in a pair of WT and *Rimbp2-/-* SGNs, showing that TTX removed action currents and revealed mEPSCs. While the amplitude of mEPSCs remained unchanged, their frequency was significantly reduced, suggesting impaired spontaneous exocytosis in *Rimbp2-/-* IHCs. Data are presented as mean ± SD (*: p<0.05, **: p<0.01). Scale bar = 25 μm. Figure 7-source data 1 Related to Figure 7c and d.

The reduced spontaneous spikes in SGNs in the *Rimbp2-/-* cochlea is likely due to reduced exocytosis from IHCs, but an alternative explanation is a change in the excitability of SGNs. We therefore injected currents in SGNs under current-clamp and studied their excitability. We found neither the input resistance (WT: 0.42±0.09 GΩ, *Rimbp2*-/-: 0.47±0.09 GΩ, p = 0.3008, Supplementary Fig. 3c), nor the resting membrane potential (RMP, WT: -62.6±1.15 mV, *Rimbp2*-/-: -62.4±1.21 mV, p = 0.7552, Supplementary Fig. 3d), nor the spike threshold (WT: -36.6±5.34 mV, *Rimbp2*-/-: -39.0±1.77 mV, p = 0.0513, Supplementary Fig. 3f) was significantly altered in the *Rimbp2-/-* SGNs, indicating that their excitability was unchanged. Furthermore, no significant difference was observed in the spike amplitude (WT: 33.0±8.12 mV, *Rimbp2*-/-: 35.5±8.47 mV, p = 0.6120, Supplementary Fig. 3e) or the spike half-width (WT: 1.49±0.32 ms, *Rimbp2*-/-: 1.76±0.25 ms, p = 0.1095, Supplementary Fig. 3g), either. We also examined the density and morphology of SGNs with immunohistochemical staining of frozen cochlear sections (Supplementary Fig. 3h), and we found no significant change between the WT and *Rimbp2-/-* cochlea. Taken together, these results suggest SGNs were not significantly impacted by *Rimbp2* deletion.

To directly address possible change in spontaneous exocytosis from IHCs, we turned back to patch-clamp recording in SGNs. Under voltage-clamp and with the large action current blocked by TTX, we recorded miniature excitatory postsynaptic currents (mEPSCs). These mEPSCs were significantly smaller in amplitude when compared to those recorded from the bouton terminals (Glowatzki E, 2002), but still detectable. We found that the amplitude of these mEPSCs was not significantly changed (WT: - 18.0±6.05 pA, *Rimbp2*-/-: -23.4±3.89 pA, p = 0.2605, Fig. 7i), its frequency was significantly reduced (WT: 2.63±0.32 events/min, *Rimbp2*-/-: 1.12±0.26 events/min, p = 0.0031, Fig. 7d). These results again demonstrated that spontaneous exocytosis from IHCs was reduced in the *Rimbp2-/-* cochlea.

### Subtle change in localization of ribbon synapses in Rimbp2-/- IHCs

Lastly, we examined the morphology and number of ribbon synapses in *Rimbp2-/-* IHCs. We collected and fixed cochleae from WT and *Rimbp2-/-* mice, and performed immunostaining with antibodies against CtBP2, to mark presynaptic ribbons, and GluR2, to label postsynaptic receptors. As shown in Fig. 8a, we found no significant difference in alignment of fluorescence puncta between CtBP2 and GluR2. We also count the numbers of puncta in IHCs, and no significant difference was found, either (Fig. 8b-d). However, when we quantified the distance of individual CtBP2 puncta to the nucleus, we found a systematic shift of their localization towards the cell’s basal pole, for all the three turns in the cochlea (Fig. 8e-g). On average, we found the shift to be 1.85, 2.81 and 2.14 μm for apical, middle and basal turn, respectively. Meanwhile, there was no significant difference in the coefficient of variation of CtBP2 localization between the WT and *Rimbp2-/-* cochlea (WT: 0.75±0.07, *Rimbp2*-/-: 0.95±0.25, p = 0.2575). In conclusion, we found that *Rimbp2* deletion caused a subtle but systematic shift of ribbon synapses towards the basal pole in IHCs.

**Figure 8.**
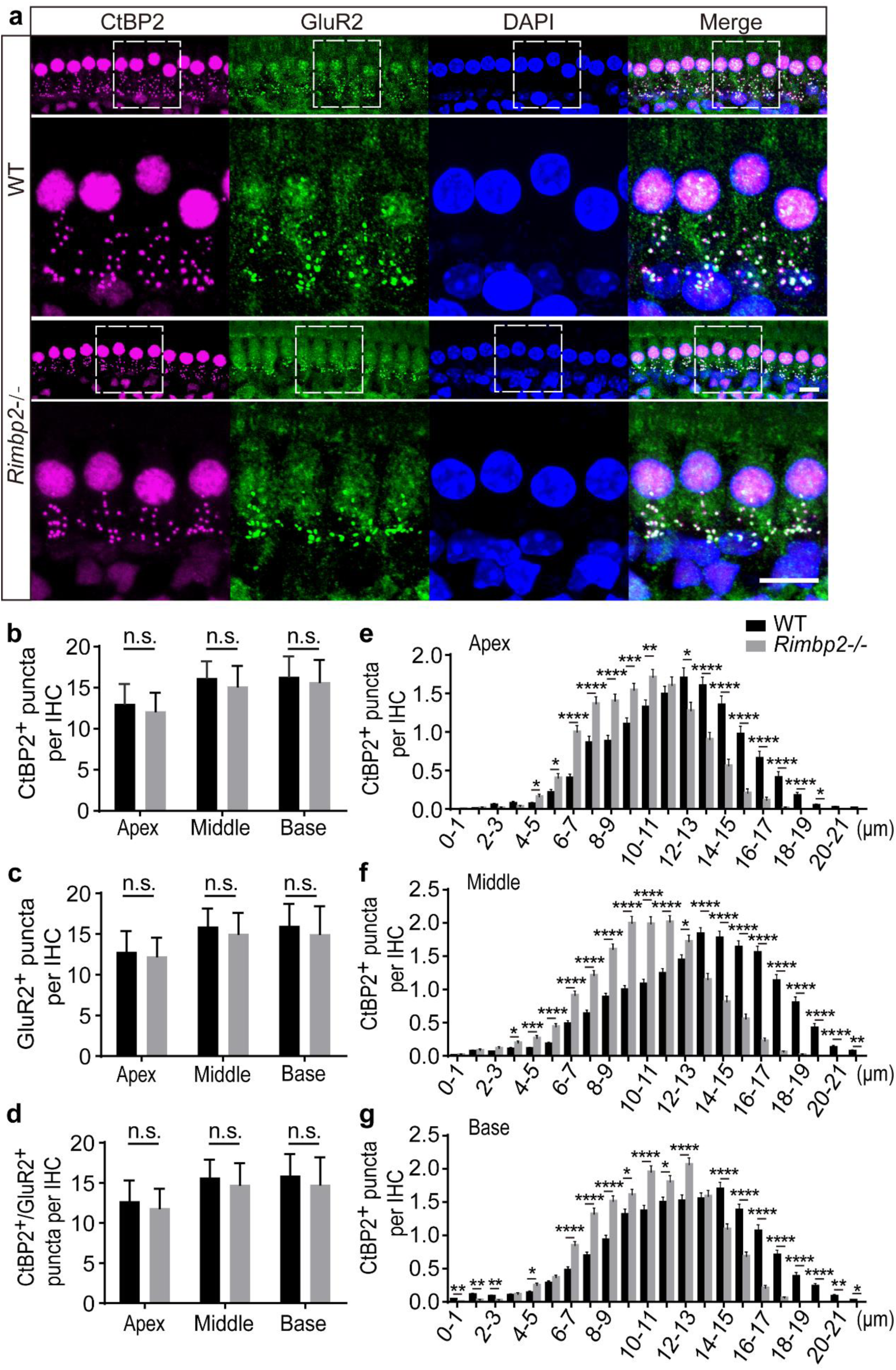
Localization of ribbon synapses in IHCs was subtly but significantly moved towards the basal pole in the *Rimbp2-/-* mouse cochlea. (a) Immunofluorescence images of ribbon synapses in WT and *Rimbp2-/-* IHCs at P21. Staining was obtained with antibodies against CtBP2 (magenta), a specific marker for synaptic ribbons, and GluR2 (green), a protein subunit for AMPA receptors. Images showed the middle turn of the cochlear basement membrane. For each panel, the area of a white dotted box is redrawn in an enlarged view below. (b-d) Numbers of immunofluorescence puncta in IHCs from the apical, middle and basal turns. There was no significant difference between WT and *Rimbp2-/-* IHCs in CtBP2 or GluR2, either counted separately or together. Data are presented as mean ± SD. (e-g) Localization of CtBP2 puncta in IHCs, measured relative to the nucleus, was subtly but significantly moved towards the basal pole. Data are presented as mean ± SEM (*: p<0.05, **: p<0.01, ***: p<0.001, ****: p<0.0001). Scale bar = 10 μm. Figure 8-source data 1 Related to Figure 8e-g.

## Discussion

In the present study, we set out to investigate the functions of *Rimbp2* in hearing. After verified the expression of *Rimbp2* in both OHCs and IHCs in the cochlea, we generated a *Rimbp2* knockout mouse model, which, in contrast with the previous model showing only mild hearing impairment (Krinner et al., 2017; Krinner et al., 2021), exhibited severe hearing loss of 20 - 50 dB (Fig. 3). Taking advantage of this new mouse model, we uncovered a multifaceted role of *Rimbp2* in promoting hearing.

### Rimbp2 and OHC survival

It is surprising that deletion of *Rimbp2*, which is widely regarded as a scaffold protein in the active zones for exocytosis (Mittelstaedt & Schoch, 2007), causes significant loss of OHCs (Fig. 4). It is not so surprising, however, given that RIMBP2 is expressed specifically and extensively in OHCs (Fig. 1). While it remains to be elucidated, the three SH3 domains in RIMBP2 could bind to membrane and/or scaffold proteins, making up the whole scaffold matrix for OHCs. Meanwhile, OHCs are subjected to constant mechanical stress in their life time (Fettiplace & Hackney, 2006), making them extremely vulnerable to mutations and/or deletions of scaffold genes (Wagner & Shin, 2019). Furthermore, in amplifying the mechanical vibrations through their electromotility (Fettiplace & Hackney, 2006), OHCs are in such a high demand of energy, requiring optimal operation of the intracellular scaffold matrix for regulating cellular metabolism (Mugabo & Lim, 2018). Many scaffold proteins, including two postsynaptic scaffold proteins, Homer1 and PSD95, are shown to be actively involved in regulating cell damage, apoptosis and cell survival (T. Chen et al., 2013; Zhou et al., 2010). Lastly, while it is not clear whether RIMBP2 is directly involved in suppressing apoptosis of OHCs and promoting their survival, at least one of its binding partners, Bassoon, and three additional presynaptic scaffold proteins, Endophilin A, Synaptojanin1 and Rab26, are shown to be linked to autophagy and related pathways in controlling cell damage and survival (Binotti et al., 2015; Okerlund et al., 2017; Soukup & Verstreken, 2017; Vanhauwaert et al., 2017).

### Rimbp2 and IHC exo cyto sis

In a previous mouse model, deletion of *Rimbp2* caused only mild hearing impairment, with hearing loss from a few dBs at high frequencies to 20 dB at low frequencies (Krinner et al., 2017; Krinner et al., 2021). Furthermore, there was only about 20% of reduction in the Wave I amplitude of ABRs. Consistently, the impact of *Rimbp2* deletion on IHC exocytosis was also mild in this mouse model, with sizeable reduction in the sustained release of SVs only, likely due to a slightly slower replenishment of SVs (Krinner et al., 2017). In the present study, we generated a new *Rimbp2* knockout mouse model, and we found severe hearing loss in the range of 20 - 50 dB, along with more than 50% of reduction in the Wave I amplitude of ABRs (Fig. 3). While the hearing loss differences between the two animal models are likely due to their different genetic backgrounds, the new animal model does lend us advantage to not only uncover the role of RIMBP2 in promoting OHC survival (see discussion above), but also examine in depth its pervasive role in regulating IHC exocytosis.

Firstly, we found that deletion of *Rimbp2* reduced RRP in IHCs by almost half (Fig. 6), suggesting RIMBP2 plays a rather significant role in priming of SVs for fusion. While this finding is consistent with studies on the Drosophila NMJ (Liu et al., 2011; Muller et al., 2015), it is in contrast with results from many other synapses (Acuna et al., 2015; Grauel et al., 2016), including a retinal ribbon synapse (Luo et al., 2017), where the size of RRP remained unchanged with *Rimbp2* deletion. In IHC ribbon synapses, similar result was reported, but upon close examination, the data actually showed an apparent trend of reduction but did not reach significance (Krinner et al., 2017). Combined this result with ours, it is clear that RIMPB2 is involved in determining the size of RRP, likely through priming of SVs. Furthermore, we found the time constant for releasing RRP was unaltered with *Rimbp2* deletion, consistent with the previous finding in IHCs that the Ca^2+^ dependency of RRP releasing was unaltered (Krinner et al., 2017), but in contrast with results from other ribbon and non-ribbon synapses where RRP releasing is often slowed or desynchronized (Acuna et al., 2015; Grauel et al., 2016; Luo et al., 2017).

Secondly, we found deletion of *Rimbp2* almost completely blocked a fast endocytosis in IHCs, likely a mechanism for observed reduction in the sustained release rate (Fig. 6). Unlike its varied effect on RRP and its release, it is almost unanimous in all synapses that deletion of *Rimbp2* significantly reduced the sustained release rate of SVs (Acuna et al., 2015; Grauel et al., 2016; Krinner et al., 2017; Luo et al., 2017). Furthermore, in all these studies, the reduction in the sustained release rate was attributed to slower replenishment of SVs. However, it is not clear what caused the slower SV replenishment: is it due to slower internal recycling of SVs or is it due to slower endocytosis? Our finding of complete blockade of the fast endocytosis with *Rimbp2* deletion provides a direct link between RIMBP2 and endocytosis, therefore an explanation for slowed SV replenishment and sustained release rate.

Thirdly, we found deletion of *Rimbp2* reduced the frequency but not the amplitude of spontaneous EPSCs recorded in SGNs (Fig. 7). This is significant because this question has not been addressed in IHCs in the previous study (Krinner et al., 2017), and because in almost all other synapses examined so far, deletion of *Rimbp2* did not alter the frequency or amplitude of spontaneous EPSCs (Acuna et al., 2015; Liu et al., 2011; Luo et al., 2017). Furthermore, spontaneous releases of SVs from IHC consist of both single and multivesicular releases (Glowatzki E, 2002; Li et al., 2009), and the fact that deletion of *Rimbp2* reduced the frequency of spontaneous EPSCs provides the first piece of evidence that *Rimbp2* is involved in multivesicular releases, a hallmark of IHC ribbon synapses. However, because our spontaneous EPSCs were recorded from the soma with significantly attenuated amplitudes, we cannot separate single versus multivesicular releases, so that it is still unclear whether the reduced release rate applies to both types of releases or specifically to multivesicular releases only.

Lastly, we found that deletion of *Rimbp2* caused a subtle but systematic change of ribbon synapse localization (Fig. 8). Specifically, we found ribbon synapses in IHCs moved around 2 - 3 μm towards the basal pole of the cell, and this localization change is parallel and systematic among ribbon synapses in that there was no significant change in the coefficient of variation of their localization (Fig. 8). This is a novel finding because no morphological change like this has been reported in any synapse yet with deletion of *Rimbp2*. While deletion of *Rimbp2* causes loosened alignment of pre- and postsynaptic density in non-ribbon synapses, it is not the case for ribbon synapses so far (Krinner et al., 2017; Luo et al., 2017). A systematic change of localization in IHC ribbon synapses is intriguing, and the underlying mechanisms remain to be further investigated.

In summary, we generated a new *Rimbp2* knockout mouse model that exhibited severe hearing loss, and we found significant loss of OHCs and pervasive change of exocytosis in IHCs, pointing to a multifaced role of *Rimbp2* in promoting hearing.

## Materials and Methods

### Animals

The *Rimbp2* knockout mice were generated with the help from Dr. Guicheng Wang, School of Life Science, Fudan University, China. The use and handling of animals were reviewed and approved by the Animal Care and Use Committee of Southeast University, Nanjing, China, in agreement with the Guide for the Care and Use of Laboratory Animals, published and updated by the National Research Council of the National Academies in USA.

### Immunohistochemistry

After animals being sacrificed, the cochleae were dissected in pre-cooled HBSS (Gibco) and fixed with freshly prepared 4% paraformaldehyde at room temperature (RT) for 1 hour. Next, the tissue was washed with PBST (0.1 M phosphate buffer, pH 7.2, with 0.01% Triton X-100) for 3 times and transferred to blocking medium containing 0.5% Triton-100 and placed at RT for 1 to 1.5 hour. Then incubated the tissue with primary antibodies (Supplementary Table 1) overnight at 4°C. After washed with PBST for 3 times, the tissue was incubated with DAPI and secondary antibodies (Supplementary Table 1) for 1 hour at RT. Lastly, the tissue was placed on a glass slide with Dako fluorescence mounting medium (DAKO, S3023), and images were taken with a Zeiss

### LSM700 confocal microscope

For preparation of frozen cochlear sections, the temporal bones were fixed with freshly prepared 4% paraformaldehyde at RT for 1 hour, and stored at 4°C overnight. After being permeabilized with 0.01% PBST for 20 minutes, the temporal bones were transferred into 15%, 20%, and 30% sucrose solutions sequentially for gradient dehydration under vacuum, and then placed at 4°C overnight. Then the temporal bones were treated with a mixture of 30% sucrose solution and OCT in a mixture ratio of 1:1, 3:7 and 3:17 sequentially, to vacuum for 1 hour each, and then placed at 4°C overnight. Lastly, they were transferred to 100% OCT, with its position adjusted so that the round window and fenestra ovali faced downwards, vacuumed for 1h, and placed at 4°C overnight. The temporal bones were vacuumed for 1 hour again the next day, quickly frozen for 20 minutes in a cryostat, and then moved to a -80°C refrigerator. Lastly, the tissues were cut into sections of 10 μm and stored at -20°C for use.

### Western Blot

After animals being sacrificed, the cochleae were dissected in pre-cooled HBSS, lysed in RIPA lysis buffer (FUDE, FD008) and cocktail (Roche, 04693132001), and then placed on ice for 10 or 15 minutes. After centrifugation at 12000 rpm, 4°C for 10 minutes, debris were removed from supernatant, and a BCA protein quantification kit (Sangon Biotech, C503061-1250) was used to measure the protein concentration. Each lysate sample was loaded to 5×SDS-PAGE sample loading buffer (Beyotime, P0015L) and boiled in 100°C water bath for 10 minutes to denature proteins. Next, proteins were separated by SDS-PAGE and transferred onto nitrocellulose membrane. Non-specific signals were blocked with 5% milk for 1h at RT, and incubated with the corresponding primary antibodies (Supplementary Table 1) at 4°C overnight. The next day, samples were washed 3 times with TBST (10 mM Tris-HCl, 150 mM NaCl, 0.1%Tween-20, pH 7.5), 15 minutes each time, incubated with the secondary antibodies (Supplementary Table 1) for 1h at RT, and then chemiluminescent agent was added to detect the signal.

### Q-PCR

Mouse cochleae were dissected and cells were lysed with Trizol-reagent (Life, 15596-018) to release nucleic acids. A cDNA synthesis kit (Thermos, K1622) was used for reverse transcription of mRNA into cDNA. Pairs of specific primers were designed to target specific DNA sequences (Supplementary Table 2). SYBR Green Master (Roche, 04913914001) was used to detect the amount of double-stranded DNA presented in the PCR system. The expression of GAPDH was used to normalize the expression levels of targeted mRNAs, and the data were analyzed with the comparative cycle threshold (ΔCt) method.

### Auditory Brainstem Responses

To record ABRs from mice, we used a TDT System III workstation (Tucker-Davis Technologies, USA), driven by the SigGen32 software running on a PC computer (Wang et al., 2017). TDT software (BioSigRZ and SigGenRZ) was used to generate stimuli to drive a speaker and record ABR signals. 10 mg/kg pentobarbital sodium was used to anesthetize the mice. Then the active electrode was placed on the top of the mice skull, and the reference and ground electrode on posterior skin of two ears to record ABR signals. Pure tone pips of 4, 8, 12, 16, 24, and 32 kHz were presented sequentially, and for each frequency sounds were presented from a sound pressure level of 90 dB in a decreasing order at 5 dB a step, until the ABR waveform disappeared.

### Patch-Clamp Electrophysiology

Whole-cell patch-clamp recording was performed in IHCs and SGNs in the apical turn of freshly excised organs of Corti, as previously described (Y. Chen et al., 2021; Sun et al., 2018). For recording in IHCs, the tissue was immersed in an oxygenated extracellular solution containing (in mM): 123 NaCl, 5.8 KCl, 5 CaCl_2_, 0.9 MgCl_2_, 10 HEPES, 5.6 D-glucose, 0.7 NaH_2_PO_4_ and 2 Na-pyruvate (290 mOsm, pH 7.40), and visualized under a 60x water-immersion objective in an up-right microscope (Olympus) (Y. Chen et al., 2021). For recording in SGNs, the external Ca^2+^ was reduced to 1.3 mM, and a K^+^-base internal solution was used for current-clamp recordings. Patch-clamp recordings were made with an EPC10/2 amplifier (HEKA Electronics, Lambrecht Pfalz, Germany), driven by a PC computer running Patchmaster (HEKA Electronics). Whole-cell membrane capacitance measurement was conducted with the lock-in feature and ‘‘Sine+DC’’ method in Patchmaster (HEKA), with sine waves of 1 kHz and 50 mV (peak-to-peak) superposed on the holding potential. All patch-clamp experiments were carried out at RT and the liquid junction potential was estimated to be 10 mV and corrected offline.

To examine exocytosis dynamics in IHCs, we plotted capacitance change (ΔC_m_) against stimulation time (t) collected from individual IHCs, and fitted the data to a combination of an exponential function for releasing of RRP of SVs (C_m,RRP_, τ_RRP_), a straight line for sustained release and replenishment of SVs (R_sustained_), and a constant to account for fast endocytosis (C_m,endocytosis_):

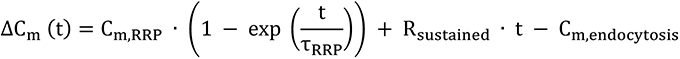

For C_m,RRP_, R_sustained_ and C_m,endocytosis_ extracted from curve fitting, the values in capacitance were converted to numbers of synaptic vesicles with a ratio of 37 aF/SV (Johnson, Franz, Knipper, & Marcotti, 2009; Lenzi, Runyeon, Crum, Ellisman, & Roberts, 1999).

### Statistical analyses

All experiments were repeated at a minimum of 3 times, data were analyzed in Microsoft Excel and Igor Pro 6.22A (WaveMetrics, USA), and the statistical tests were performed in GraphPad Prism 6. Statistical significance was evaluated with a two-tailed, unpaired Student’s t-test. p<0.05 was considered statistically significant. All data are presented as mean ± SD in both text and figures (except figure 8 e-g). The final form of figures was composed in Adobe Illustrator CS6 software.

## Additional information

### Author Contributions

M. Liao performed most experiments in immunofluorescent staining, analyzed data and drafted the manuscript; X. Chen completed patch-clamp recordings on SGNs and analyzed the data; L. Lu performed part of immunofluorescent staining and acquired funding; R. Guo provided guidance in experimental designs; P. Zhang and Y. Li completed patch-clamp recording in IHCs and analyzed the data; Y. Zhang and Q. Fang helped in experimental troubleshooting; Y. Hu helped with generation of *Rimbp2-/-* mice; J. Chai and X. Chen assisted in data analysis; M. Tang, X. Gao and S. He provided technical and experimental advice; H. Li, G. Li and R. Chai conceived the study, designed experiments, assumed supervision, acquired funding and edited the manuscript.

---

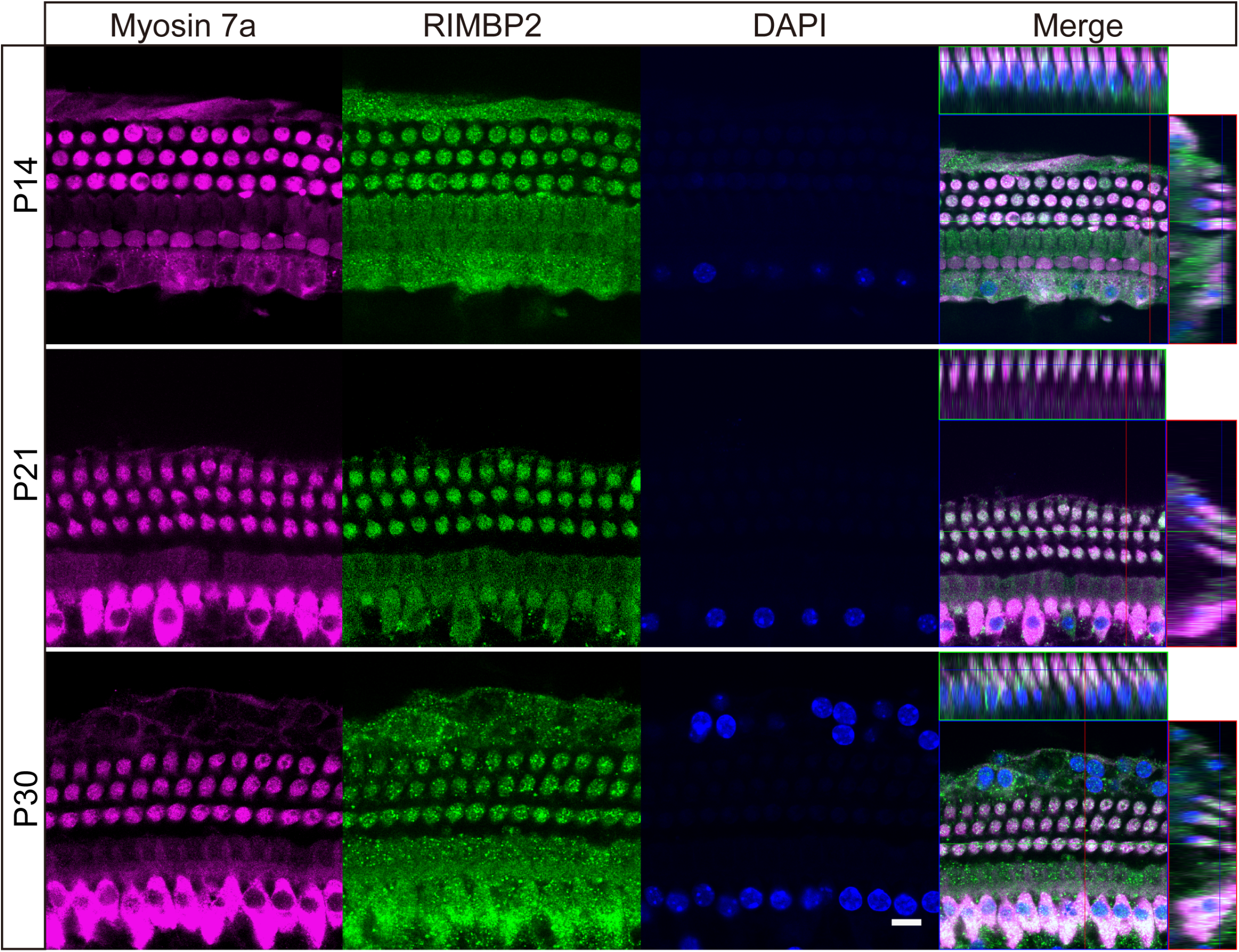

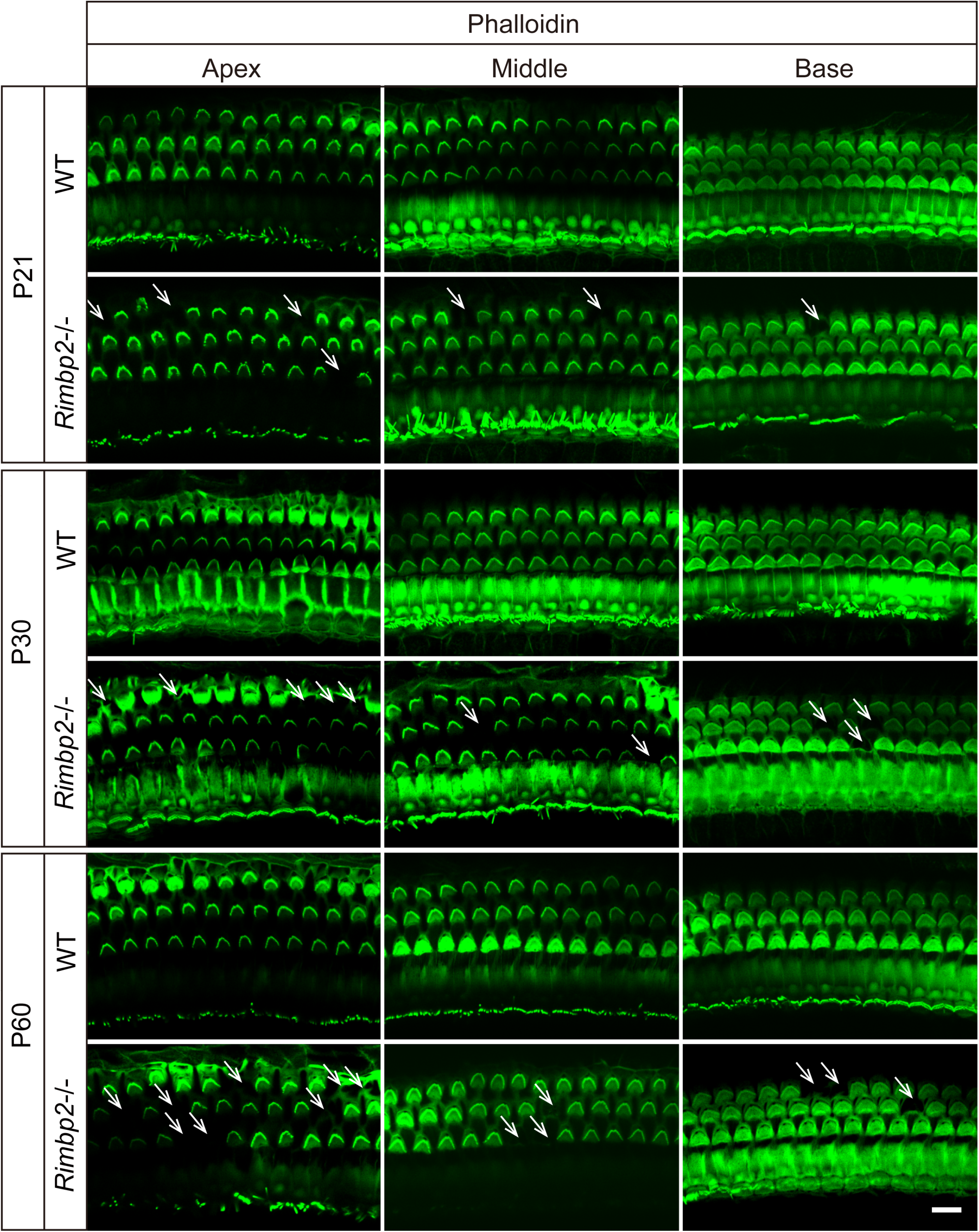

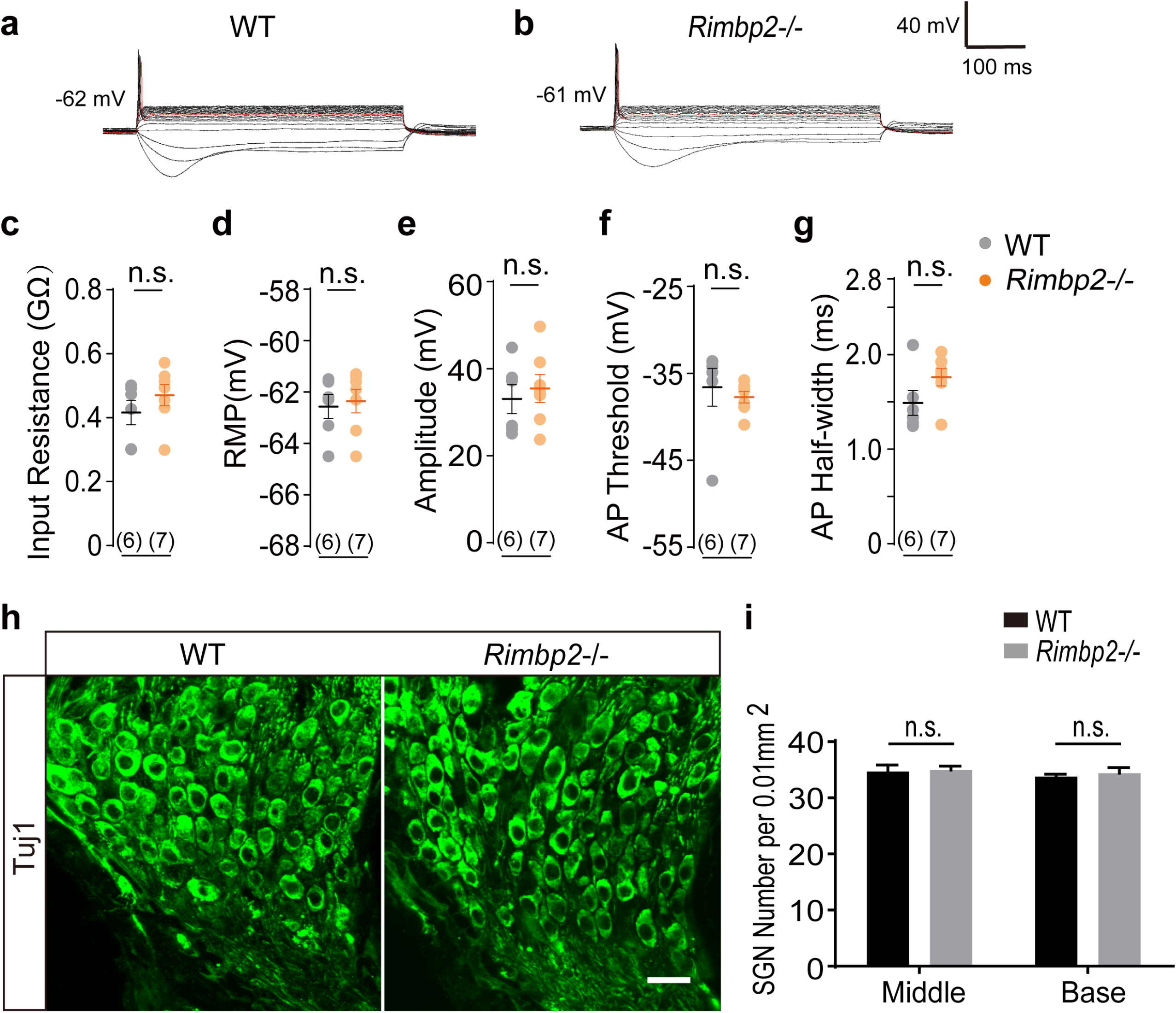

